# Effects of context changes on memory reactivation

**DOI:** 10.1101/2024.03.20.585920

**Authors:** Şahcan Özdemir, Yağmur Damla Şentürk, Nursima Ünver, Can Demircan, Christian N.L. Olivers, Tobias Egner, Eren Günseli

**Affiliations:** Department of Psychology, Sabancı University, Istanbul, Turkey; Department of Experimental and Applied Psychology, Vrije Universiteit, Amsterdam, Netherlands; Department of Psychology & Neuroscience, Duke University, Durham, NC, USA

## Abstract

While the influence of context on long-term memory (LTM) is well-documented, its effects on the interaction between working memory (WM) and LTM remain less understood. In this study, we explored these interactions using a delayed match-to-sample task, where participants (6 Male, 16 Female) encountered the same target object across six consecutive trials, facilitating the transition from WM to LTM. During half of these target repetitions, the background color changed. We measured the WM storage of the target using the contralateral delay activity (CDA) in electroencephalography (EEG). Our results reveal that task-irrelevant context changes trigger the reactivation of long-term memories in WM. This reactivation may be attributed to content-context binding in WM and hippocampal pattern separation.

**Significance Statement:** Understanding the mechanisms of memory updating in response to changing contexts is vital because context plays a pivotal role in shaping long-term memories. This study demonstrates, for the first time, that an irrelevant context change triggers the reactivation of learned memories in visual working memory. This observation underscores the importance of multi-memory interactions during context updating. Challenging traditional memory models that postulate mandatory reactivation of long-term memories upon each use, our results instead reveal a selective reactivation process, especially during transitions to new environments. This finding elucidates the adaptive nature of memories and enhances our understanding of memory storage and retrieval processes.

## Introduction

Working memory (WM) is critical in storing novel task rules and their associated information. When used repeatedly, these representations transfer to long-term memory (LTM; Carlisle et al., 2011; Gunseli et al., 2014ab; Logan, 1998). LTM is strongly intertwined with the context in which memories are formed and utilized. For example, retrieval is more successful when it shares the same context with encoding (Godden & Baddeley, 1975; Robin et al., 2016; Tulving & Thomson, 1973). Changes in context also determine the temporal structure of episodic memories by generating event boundaries at context transitions (e.g., Clewett et al., 2017; Wang et al., 2022; Güler et al., 2023; Güler et al., 2024; Nolden et al., 2024). These findings suggest that context strongly influences how an item is preserved in LTM.

Although the effects of context change on LTM encoding and retrieval are well established, its influence on the interaction between WM and LTM remains unclear. For instance, when people are required to switch between different task rules, the associated rule-updating process results in a reactivation of task-relevant information in WM (Şentürk et al., 2024). By contrast, it is not currently known whether a change in context would result in a similar WM updating process as a change in task rule. There are several reasons why context changes may have this effect on WM. First, reactivating information in WM upon a context change may be critical for individuating similar memories from different contexts, possibly by associating each memory with a unique context or via hippocampal pattern separation. Hippocampal pattern separation enhances the individuation of memories via dissociating activity patterns of similar memories (Amer & Davachi, 2023; O’Reilly & McClelland, 1994; Yassa & Stark, 2011). This process can occur through cortical activity (Wimber et al., 2015), which is a feature of WM storage (Vogel & Machizawa, 2004; Thrower et al., 2023; Fuster & Alexander, 1971; Funahashi et al., 1993). Second, reactivating memories upon context switches could be one of the contributing factors to the segmented nature of episodic memories (Güler et al., 2023; Sols et al., 2017). Given this intimate relationship between context, task-relevant items, and task rules, we hypothesized that context switches, akin to task switches, should trigger the reactivation of long-term memories in WM, to adjust to the novel context.

To investigate how context change affects the interplay between WM and LTM, we employed a measure of memory reactivation and assessed how it responded to contextual changes. Participants performed a recognition task in which the target object was repeated across trials. During this repetition, a task-irrelevant context (the background color) occasionally changed. We used the contralateral delay activity (CDA), which has become a widely used EEG index of active WM storage (Luria et al., 2016; Vogel et al., 2005). This first of all allowed us to assess learning and transfer to LTM, as this has been shown to be accompanied by a decline in CDA with increasing repetition of the same item across trials, (Carlisle et al., 2011; Gunseli et al., 2014ab, Grubert et al., 2016; Reinhart et al., 2014; Reinhart & Woodman, 2014; Reinhart et al., 2016; Şentürk et al., 2024). Second, and most important, the CDA allowed us to test our main hypothesis: Considering that LTM is reactivated in WM to adapt to changes in the environment (Artuso & Palladino, 2011; Cai 2020; Şentürk et al., 2024) and that adaptation processes like hippocampal pattern separation may occur via cortical reactivation associated with WM (Wimber et al, 2015), we hypothesized that context shifts would trigger the reactivation of information handed off to LTM back in WM. If so, this should become evident as a rebound in the CDA for the already-learned item. In contrast, if WM is not context-sensitive, then information handed off to LTM should remain in LTM without being reactivated in WM despite contextual shifts.

## Methods

### Participants

29 students from Sabancı University participated for course credits in this study, and 7 participants were excluded from further analysis due to exclusion criteria (explained below). Data collection was conducted following our lab’s guidelines to minimize COVID-19 transmission risk (Hasşerbetçi & Günseli, 2019). The analyses were conducted with 22 participants (Meanage = 21.7, SD = 2.9), 6 males and 16 females. We determined the target participant number based on our previous work that used a similar experimental design (Şentürk et al., 2024). Effect sizes from four studies that used CDA as a memory load measure were averaged (Berggren & Eimer, 2016; Gunseli et al., 2014b; Reinhart et al., 2016; Xie & Zhang, 2018). Following the guidelines by Schönbrodt and Wagenmakers (2018) and Dienes (2021), and the associated R package by Schönbrodt & Stefan (2019) (github.com/nicebread/BFDA), a sequential design approach was conducted. Cauchy distribution as an uninformed and objective distribution was chosen as prior with half of the estimated effect size as the scaling parameter (Dienes, 2021). We calculated the minimum number of participants as 20 by estimating the number of participants needed for a power of .90, and a false positive rate of .02 under the estimated effect size d=1.12. The number of participants as a stopping rule was estimated as 70 with the smallest effect size in the abovementioned studies, which is d= 0.48. Bayes Factor was planned to be calculated first when the number of participants reaches 20 then aimed to be calculated after every 5 new participants.

Data collection was decided to continue until the Bayes Factor is between 1/6 and 6 or the number of participants reaches 70. We collected 22 participants in total after exclusion. Because we were late in checking the statistical analyses of 20 participants, we exceeded the specified number of 20 without calculating the Bayes Factor. We realized the mistake with 22 participants. We first calculated the BF for the first 20 participants and the result was above 6, then we calculated the BF again with all 22 participants and the results were still similar (above 6) so we stopped the data collection at 22 participants.

Since color served as context in this study, before the experiment, each participant took an online Ishihara Color Blindness test, which is suggested to be as efficient as traditional paper tests (Marey et al., 2015). Participants viewed plates, which consist of solid circles of various colors and sizes that form a number or a pattern. They indicated the number written on the plate or counted the number of lines on the pattern. If they respond correctly to less than 13 plates out of 38, they are excluded from the experiment due to red-green color vision deficiency (Marey et al., 2015).

After the EEG data artifact rejection process (see below), participants who had less than 60 non-rejected trials per condition or with accuracies lower than 80% were excluded from further analysis. New participants were recruited to replace excluded participants. This study was approved by the Sabancı University Ethical Committee. All participants signed a consent form to participate in the study.

### Ethics Statement

This study was performed in line with the principles of the Declaration of Helsinki. The ethics approval was granted by the Sabancı University Research Ethics Committee.

### Stimuli

Images that were used in the current study (Şentürk et al., 2024) were brought together with additionally collected images to create a set of 2880 images of real-world objects (Konkle et al., 2010; Konkle & Caramazza, 2013; Konkle & Oliva, 2012; Google Images) in total. Images were resized to contain approximately the same number of non-transparent pixels. Half of the images were selected as target objects and the other half were selected as non-target. Target objects were separated into two groups, animate and inanimate. Animate and inanimate groups contained 60 object categories, each category consisting of 12 pictures, thus making 1440 target objects in total. Each object (7.1° x 7.1°) was presented once as a target throughout the experiment. Each category became a target twice. Two objects from each object category were used as targets during the experiment.

Each trial was presented with a color as a context. Context color could be either red (HSL: 228, 44, 146) or green (H:71, S:67, L:145). The context color was presented as a rectangular frame surrounding the center of the screen (16.7° x 10°). The experiment had 1440 trials. Each target object was repeated for 6 trials in a row. Through these repetition series, the context could change in the 1st and/or 5th repetition. Participants viewed the experiment 85 cm away from the computer screen. The background color of the experiment was grey. The location cue was a vertically halved, bicolored circle (0.35° x 0.35°), with one side being navy blue (H:240, S:100, L:25) and the other side orange (H:38,5, S:100, L:50). For a given participant, a particular color indicated the side of the screen on which the target was presented, and it was counterbalanced across participants.

### Design and Procedure

#### Trial design

The trial design is depicted in Figure 1. The task was to indicate if the probe item matches the target item. Each trial started with a presentation of the context color for 1250 ms. Then, the location cue appeared for a jittered duration of 800-1400 ms and remained on the screen until the probe was at the center of the screen. Following the location cue, two objects were presented. One of the objects was presented at the left and one to the right of the location cue, for 500 ms. Participants were instructed to memorize the target presented on the cued side. They were instructed to fixate on the location cue. Next, there was a retention interval of 900 ms during which the cue remained on the screen. After the retention interval, the probe was shown at the center of the screen. On each side of the probe, there were two response labels, “Same” and “Different” (0.7°). Participants pressed the left or right arrow key on a Turkish QWERTY computer keyboard to indicate their response (Figure 1; e.g., the left arrow to respond Same and the right arrow to respond Different). The locations of each response label were counterbalanced across participants. The probe remained on the screen until response or up to 1600 ms. Participants received visual feedback upon their response (‘correct’ or ‘incorrect’) or after 1600 ms (‘miss’). The feedback was shown at the center of the screen for 300 ms (5° x 5°). Lastly, a blank inter-trial interval was jittered between 300 and 700 ms. The context color frame was presented in the background (16.7° x 10°) throughout the trial except the inter-trial interval.

**Figure 1.**
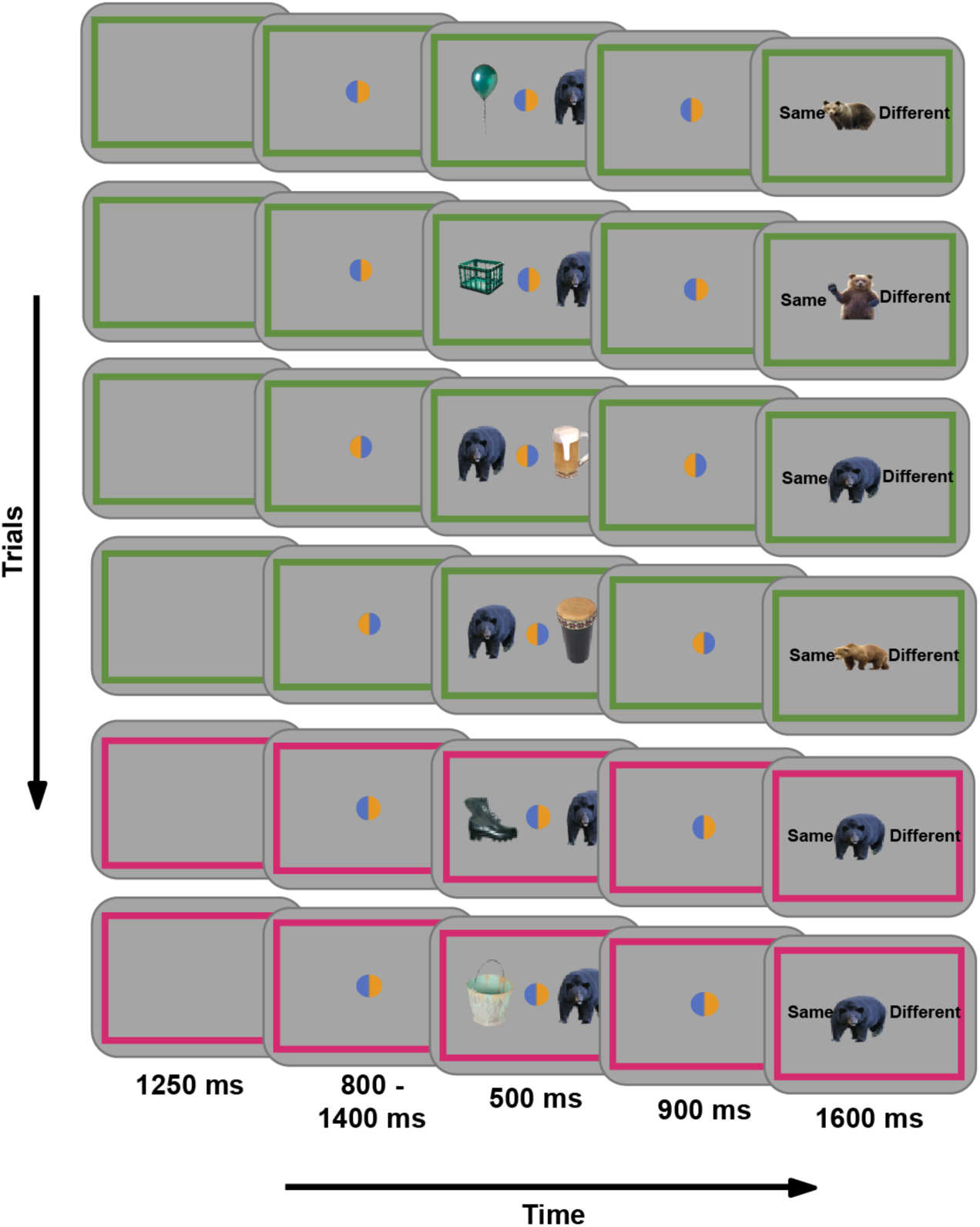
Illustration of the experimental procedure. Each trial will start with the presentation of context, i.e., the background color, which will be filled in the real experiment but shown here only as a frame for illustration purposes. Then, participants will see the location cue that indicates which object will be the target object. Following the location cue, the target object was shown at the indicated side and the non-target object was at the contralateral side. After a brief retention interval, participants will be shown a probe object and will indicate if it is the same as the target.

#### Trial distribution & block design

On each trial, the probe and the target were from the same object category. For each condition, the target was equally likely to be on the left or right on the target display. The target type was equally likely to be animate or inanimate. The experiment began with a practice session of 25 trials minimum. The objects used during practice weren’t shown in the main experiment. The practice session was repeated until participants achieved at least 80% accuracy. The experimental session was divided into 38 to 42 blocks of approximately 40 trials. The block and trial numbers varied to ensure each block starts with a new context and a new target. At the end of each block, participants were informed about their accuracy and were able to take a self-paced break.

### Behavioral analysis

Our main hypotheses concern the EEG data and are not directly related to the behavioral performance pattern. However, as in previous studies using this protocol, we expected to observe a decrease in reaction time with target repetition (Carlisle et al., 2011; Gunseli et al., 2014a). To test this, we performed a Bayesian paired sample t-test to compare reaction times of 1st and 5th target repetitions in old context trials.

### EEG recording and registered analysis

The electroencephalogram (EEG) was recorded from 32 sintered - AG/AgCl electrodes. The electrodes were positioned at International 10/20 System sites and mounted in an elastic cap using Brain Products actiCHamp (actiCHamp Plus, Brain Products GmbH, Gilching, Germany). The vertical EOG electrodes were located at 2 cm above and below the right eye, and the horizontal EOG electrodes were located at 1 cm lateral to the external canthi. We performed the analysis of the EEG data using MATLAB (2022), the EEGLAB toolbox (Delorme & Makeig, 2004), and custom code. Trials containing ocular artifacts that are recorded by EOG, and EEG noise such as blocking, muscle noise, saturation, etc., were detected manually via visual inspection. Trials with such artifacts were excluded from further analysis. We applied a filter to EEG data by IIR Butterworth filter with a band-pass of 0.1-40 Hz using the *pop_eegfiltnew.m* function of EEGLAB. The online reference electrode was located in the right mastoid, and then the data was re-referenced offline to the average of left and right mastoids. A baseline period of 200 ms prior to stimulus onset was included in the ERP analysis.

We used the CDA amplitude as a validated tool to assess WM storage based on four lines of findings. First, previous research showed that CDA scales with the number of items kept in WM and reaches an asymptote at individuals’ WM capacity (Luria et al., 2016; Vogel & Machizawa, 2004; Vogel et al., 2005). Second, the CDA has been shown to reflect the number of items in WM and not their complexity (Ikkai et al., 2010). Third, the CDA decreases when the same item is repeatedly stored in WM, reflecting the handoff of the item’s representation from WM to LTM (Carlisle et al., 2011; Gunseli et al., 2014ab, Grubert et al., 2016; Reinhart et al., 2014; Reinhart & Woodman, 2014; Reinhart et al., 2016; Şentürk et al., 2024). Thus, it is a suitable tool to measure WM involvement in storing items. Fourth and last, the CDA was found to increase following task rule changes (Şentürk et al., 2024) and cues that signal high reward (Reinhart & Woodman, 2014), deeming it possible to observe a similar increase following context change.

The CDA was computed at P7/8, PO3/4, PO7/8, and O1/2 electrodes (Günseli et al., 2019; Ikkai et al., 2010; Vogel & Machizawa, 2004; Vogel et al., 2005) by subtracting the activity of ipsilateral channels from the contralateral ones relative to the target position between 400 to 1400 ms after target onset (retention interval). To test if new targets are stored in WM (Carlisle et al., 2011; Gunseli et al., 2014a,b; Reinhart et al., 2016), the CDA in new target trials were compared against zero using two one-sample t-tests, one for new contexts and the other for old contexts.

Next, to test if repeated targets are being handed off from WM to LTM (Carlisle et al., 2011; Gunseli et al., 2014ab, Grubert et al., 2016; Reinhart et al., 2014; Reinhart & Woodman, 2014; Reinhart et al., 2016; Şentürk et al., 2024), we compared the 1st and the 5th target repetitions using a Bayesian paired-samples t-test. For this analysis, we used old context trials only. To assess if this difference showed a linear or a quadratic CDA trend through a target repetition series (Figure 2), a trend analysis with repeated measures ANOVA was conducted. Last and importantly, to assess whether a context change results in the targets stored in LTM being reactivated in WM, we used a Bayesian paired samples t-test to compare the CDA in new vs. old context trials only for the 5th target repetition where a handoff to LTM is expected based on previous literature (Carlisle et al., 2011; Gunseli et al., 2014 a,b; Reinhart et al., 2016).

**Figure 2.**
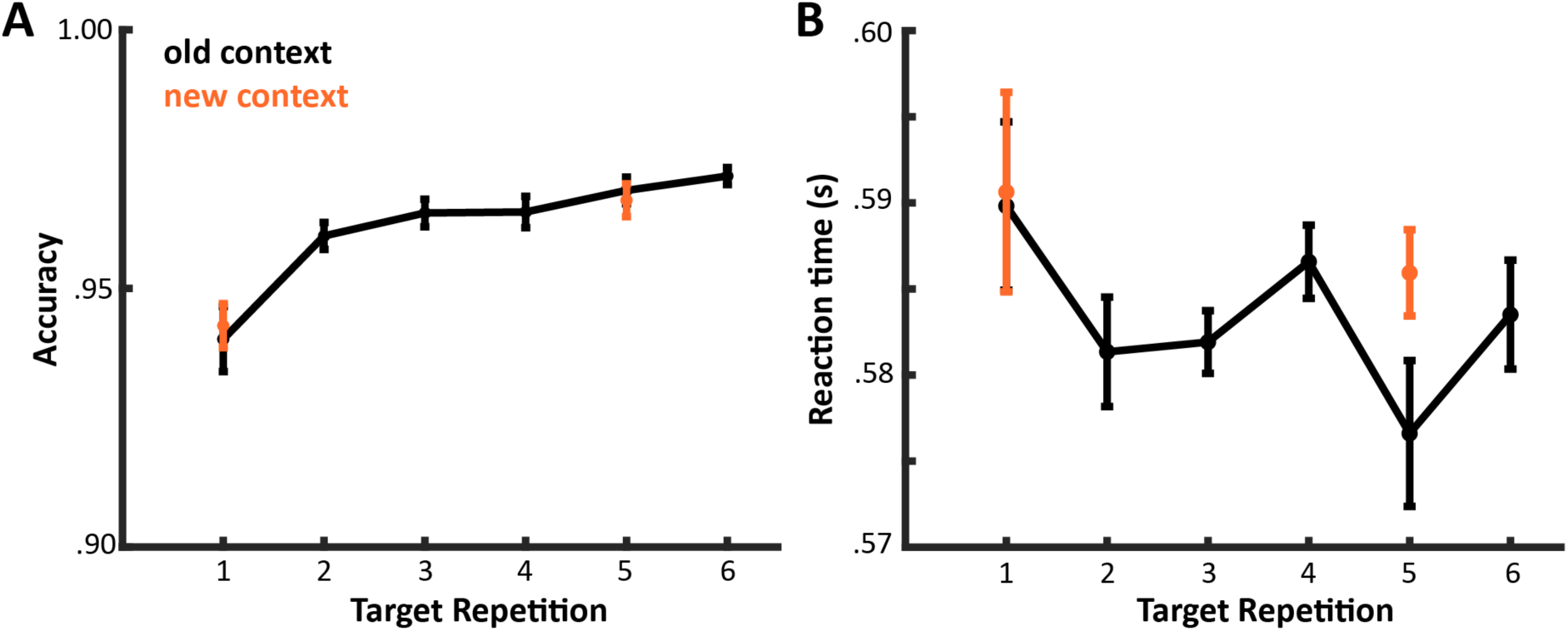
Behavioral results. (A) The accuracy increased with target repetition. (B) The reaction time remained unchanged across target repetitions. Neither were affected by context changes.

### Exploratory analysis

We calculated the N2pc, which is an index of attentional selection (Eimer, 1996; Hickey et al., 2009). Similar to CDA, N2pc was calculated as a difference between contralateral and ipsilateral activity regarding the target position, time-locked to the onset of the memory item. This difference was calculated between 250 ms and 350 ms based on visual exploration of our data and our previous study with a similar design (Şentürk et al., 2024). Time-frequency analysis was conducted to explore the involvement of attentional processes in memory reactivation (Fukuda & Woodman, 2017; Günseli et al., 2019). Bilateral and contralateral alpha-band suppression were analyzed in the time course between 500-1200 ms. The same channels with the CDA analysis (P7/8, PO7/8, and O1/2) were used in the calculation. We analyzed the power of frequencies between 4 and 50 Hz to see if the effects we observed were alpha band specific or propagated to other frequencies. To calculate bilateral alpha-band suppression, we defined our frequencies between 8-12 Hz (Günseli et al., 2019; Woodman et al., 2012) on a logarithmic scale. For each frequency, a sinusoid (ei2ft) was created then these sinusoids were converted to Morlet wavelets by being tapered with a Gaussian (n (e-t2/2s2; s is the width of the Gaussian; s = /(2f); denotes for number of cycles created for wavelet). We padded zero to the beginning and the end of our data as half of the length of our Morlet wavelets. Our epoched data were rearranged as one continuous EEG data. Fast Fourier Transform (FFT) was applied to both the EEG and Morlet waves. The dot product of the Fourier-transformed EEG data and Fourier-transformed Morlet wavelet was calculated for each frequency. Then inverse FFT was applied to each dot product. With this procedure, the EEG data became convoluted for each Morlet wavelet. Then, we performed baseline normalization and decibel (dB) conversion. Each baseline was calculated by averaging the power activity between 500-200 ms before the memory item onset of all trials. The power activity in each trial was divided by this baseline and converted to dB. Then, for the analysis regarding the alpha band suppression, we averaged the dB values between 8-12 Hz, and 400-1200 ms, over trials for each condition. We have chosen a slightly shorter time window compared to the CDA calculation to prevent probe-related activity from contaminating the power due to the temporal smearing caused by convolution.

A similar procedure was applied to calculate the contralateral alpha suppression. However, for contralateral alpha suppression, the baseline was calculated between 500-200 ms before the location cue rather than the memory onset. Since the location cue was given before the memory representation, lateralization can be expected in the baseline time window that reflects the expected attention (Ikkai et al., 2016). This might result in the cancellation of lateralization in the retention interval due to baseline removal with a lateralized baseline. This difference in baseline time range was applied to get a clear baseline without lateralization. This baseline normalization was applied to each condition separately, unlike the bilateral time-frequency decomposition. Then the power values from the selected channels (P7/8, PO7/8, and O1/2) that are contralateral to the target item position were subtracted from the ipsilateral channels (Günseli et al., 2019).

## Results

### Behavioral Results

#### 2.3.1.1 Reaction times

There was anecdotal evidence for no difference in reaction times across target repetition 1 and target repetition 5 (old context TR1 vs old context TR5, BF10 = 0.669; t(21) = 1.597, *p* = .125). Moreover, there was also anecdotal evidence for equal reaction times at the new and old context trials’ first repetitions (BF10=0.225; t(21)= 0.132, *p*=0.896). Also, there was anecdotal evidence for an increase in RT with a change in context at the 5th repetition (BF10 = 1.536; t(21) = 2.164, p = .042). When we conducted a trend analysis with repeated measures ANOVA, repetition showed no linear contrast (t(105) = -1.357; *p* = .178). These results suggest that RT was mostly unaffected by target repetitions and contextual shifts.

### Accuracy

We found strong evidence for an increase in accuracy with repetition (old context TR1 vs old context TR5, BF10 = 91.247; t(21)=-4.265, *p* < .001; linear contrast with repetition, t(105)=5.923, p < .001). Also, there was anecdotal evidence for equal accuracy at the 5th repetition between change in context (M=.96, SD=.02) and old context (M=.96, SD=.02; BF10= 0.243; t(21) =0.432, *p* = .67). Thus, while accuracy improved across target repetitions, it was mostly unaffected by context shifts.

### EEG Results

#### Contralateral delay activity

We compared the CDA amplitudes of the 1st repetition trials against 0 to test whether there is a CDA when a novel object is presented. As depicted in Figure 3A, the 1st repetition trials’ CDA values were more negative than 0 (old context, BF10=34470; t=-7.134, *p* < .001; new context, BF10=43375; t(21)=-7.252, *p* < .001). Then we compared the 1st and 5th repetitions of old context trials to test the transition of the item from WM to LTM. Results showed strong evidence that the CDA amplitude of the 5th repetition was different from the 1st (BF10=2406; t(21)=-5.815, *p* < .001), which suggests the representation of the target item was transferred to LTM with repetition of the same task and item (linear trend, t(105) =6.859, *p* < .001).

**Figure 3.**
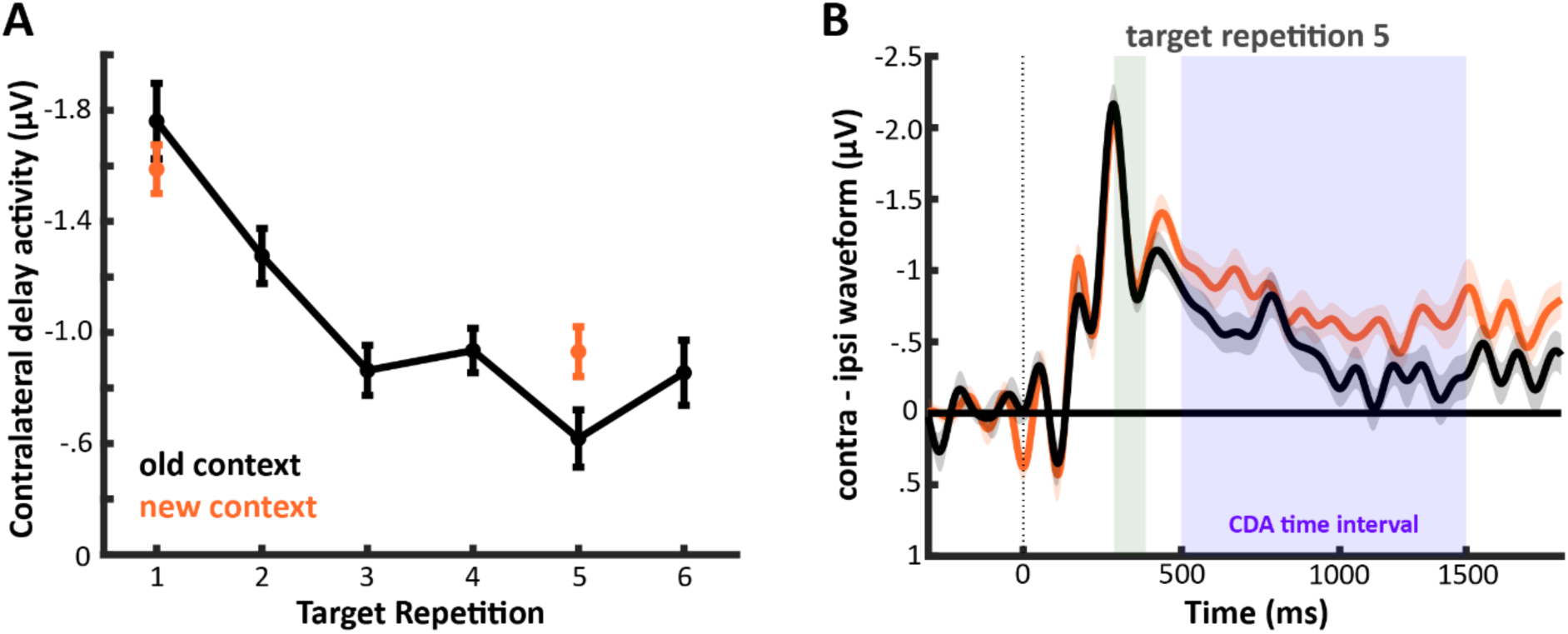
Contralateral delay activity. A) CDA changes through repetitions. CDA amplitude decreased with target repetition. CDA was recovered at the 5th repetition when there was a contextual change. B) Contralateral - ipsilateral waveforms relative to the target side for repeated and switched context trials at the fifth target repetition. The shaded areas refer to the time windows for N2pc and CDA. The recovery with a context change was present for the CDA but not the N2pc. The error bars represent the standard error of the mean for the context repeat vs switch condition difference.

Then, we compared the 5th repetition trial regarding whether there was a change in context or not. When there was a change in background color, we observed strong evidence for a recovery in the CDA (BF10=8.858; t(21)=-3.131, *p* = .005; Figure 3B). We conclude that a change in context triggers the reactivation of the task-relevant item in WM. Lastly, we tested if a similar increase is present for novel targets. The CDA for novel targets did not differ for old context and new context trials (BF10 = 0.326). This finding suggests that a CDA increase via a contextual shift at target repetition 5 is unlikely to represent a non-reactivation-related cognitive boost, as a similar shift should have been present for contextual shifts at target repetition 1.

#### N2pc

For a new target, there was strong evidence for an N2pc (BF10=767.442, t(21)=-5.269, *p* < .001) suggesting that the memory item was attended at the memory display. N2pc didn’t show any changes regarding the contextual changes. However, with the first repetition, we observed a strong increase in N2pc (BF10=55.982; t(21)=4.033, *p* < .001) (Figure 3A). After this increase, it didn’t show any changes due to context changes (BF10= 0.240; t(21)=0.401, *p* = .692).

#### Alpha-band suppression

The alpha-band suppression provided a pattern similar to the CDA (Figure 4C). For a new target, there was strong evidence for a bilateral alpha-band suppression (BF10 = 5899.886, t(21) = -6.250, *p* < .001). Moreover, there was a decrease with the repetition of the same target (old context TR1 vs old context TR5, BF10=396.46; t(21)=-4.957, *p* < .001), which was further supported by the linear trend observed with repetition (ANOVA linear contrast, t(105)=8.023, *p* < .001). Lastly, the comparison between old-context and new-context trials at the 5th repetition showed that the alpha-band suppression recovered (BF10 = 1830; t(21) = -5.683, *p* < .001). These results suggested that overall attentional resources declined with repetition and recovered with a contextual change (Reinhart & Woodman, 2014; Şentürk et al., 2024). This increase in alpha suppression could reflect memory reactivation of the task-relevant item or the novel context (Fukuda & Woodman, 2017) given that the global alpha suppression explored here is not specific to the hemisphere that represents the memory item.

**Figure 4.**
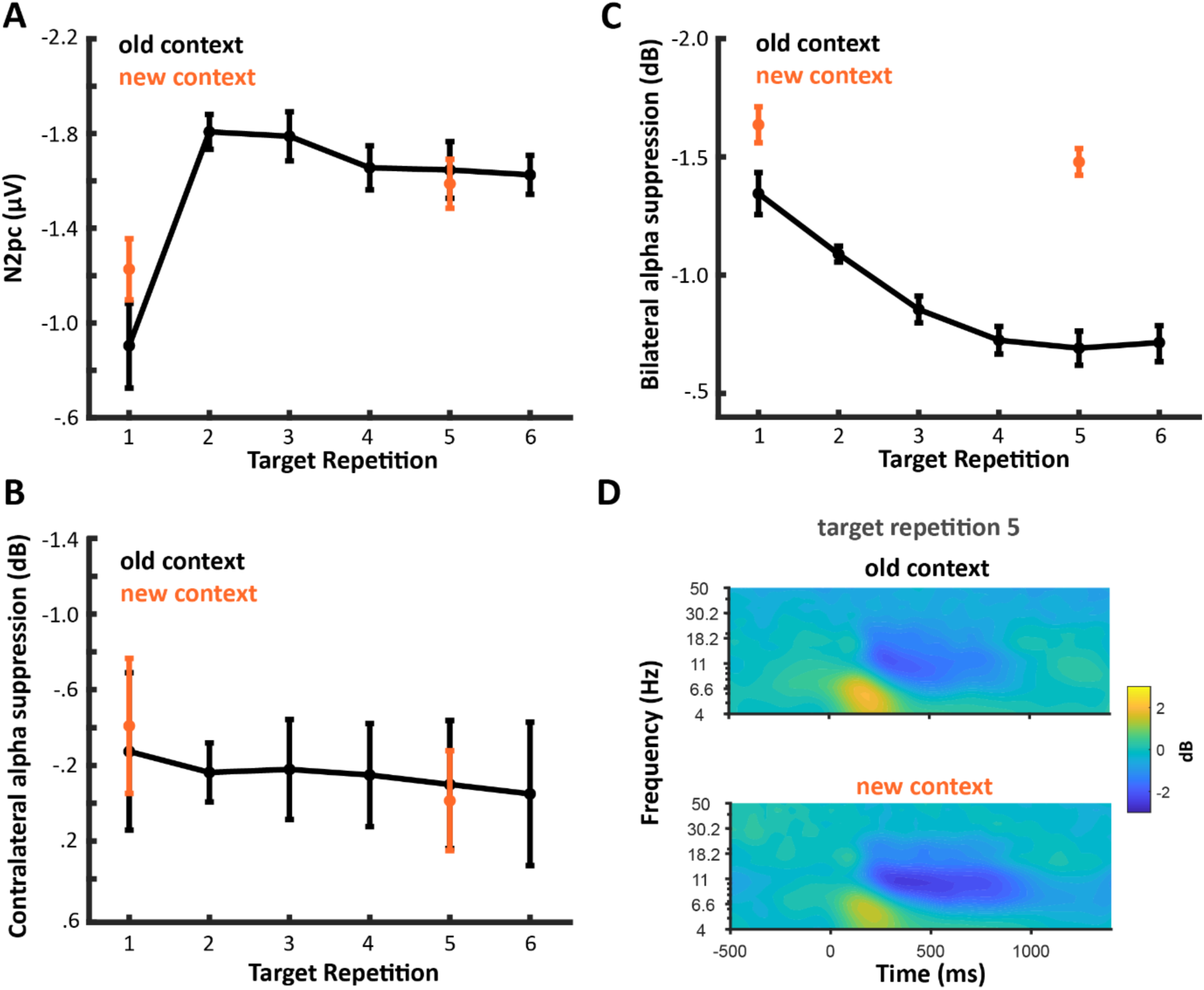
Exploratory analysis. A) Changes in N2pc through repetitions. N2pc showed a strong increase with the first repetition but responded neither to repetition nor context changes. B) Contralateral Alpha Suppression through repetitions. Contralateral Alpha Suppression was stable across repetitions and context changes. C) Bilateral Alpha Suppression through repetitions. Bilateral Alpha Suppression decreased with target repetition and increased with a context change at the 5th target repetition. D) Broadband (4-50 Hz) power at target repetition 5, separately for new vs. old context conditions.

#### Contralateral alpha-band suppression

We analyzed the spatial lateralization in the alpha-band suppression regarding the object position (Figure 4E). For a new target, there was strong evidence for a contralateral alpha-band suppression (BF10 = 29.585, t(21) = -3.727, *p* = .001) suggesting that the memory item was attended during its storage within WM. Alpha band suppression showed moderate evidence for no change regarding the repetition of the target or contextual changes (BF10 = 0.223, t(21) = .048, *p* = .96). This result suggests that participants continued to attend to the memory item’s location across repetitions. This contrast between CDA and lateral alpha is in line with our previous findings and highlights a dissociation between selective attention and storage within WM (Günseli et al., 2019; Hakim et al., 2019; Şentürk et al., 2024; van Driel et al., 2017).

#### EEG-Behavioral relations

We examined whether the reactivation in WM predicts any behavioral performance of the participants. To achieve this, we subtracted the CDA amplitudes at the 5^th^ repetition old-context trials from the 5th repetition new-context trials. Similarly, we subtracted the reaction times and accuracies at the 5th repetition of old-context trials from the 5th repetition of new-context trials. Then we checked whether the changes in CDA due to contextual changes predict any behavioral performance by analyzing correlations. CDA differences with contextual changes predicted neither a difference in accuracy nor in reaction times (CDA-RT, Pearson’s r = -0.267, *p* = .23, CDA-Accuracy, Pearson’s r = -0.087, *p* = .701).

## Discussion

We examined whether task-irrelevant context changes cause memory reactivation. Our EEG index of WM, the CDA, diminished for repeated targets, replicating previous work on the transfer from WM to LTM (Carlisle et al., 2011; Gunseli et al., 2014ab, Grubert et al., 2016; Reinhart & Woodman, 2014; Alfandari et al., 2019; Şentürk et al., 2024). Importantly, with a change in context, the CDA re-emerged, indicating that task-irrelevant context changes elicited the reactivation of task-relevant information in WM.

WM minimizes processing costs whenever possible (Chota et al., 2023; Draschkow et al., 2021; de Jong, Wilhelm, & Akyürek, 2023; Mızrak & Oberauer, 2021; Stokes, 2015; Yücel et al., 2023). In our experiment, memory reactivation was not associated with a behavioral benefit but only with a metabolic cost, reflected in increased CDA. This metabolic cost without behavioral benefits suggests that memory reactivation is not strategic but involuntary.

### Why are memories reactivated following context changes?

Reinhart and colleagues found that memories are reactivated when participants were instructed to improve performance (Reinhart & Woodman, 2014) or signaled high reward (Reinhart et al., 2016), suggesting high stakes facilitate memory reactivation. Mızrak and Oberauer (2021) suggested reactivation is reserved for memories that partially overlap with former memories, aligning with WM’s role in resolving proactive interference (Engle, 2002). We recently demonstrated that LTMs are reactivated in WM when there is a change in task rules. In the present study, we show that changes in task-irrelevant information also evoke memory reactivation. Together with Şentürk et al. (2024), our findings suggest memory reactivation helps adapt to *novel* settings, irrespective of stakes or proactive interference.

We propose two adaptive functions of memory reactivation. The first is pattern separation. The hippocampus separates similar events to prevent interference (Amer & Davachi, 2023; O’Reilly & McClelland, 1994; Yassa & Stark, 2011). Encountering the same item in a different context might signal it as distinct, encouraging pattern separation, which could benefit from cortical reinstatement associated with WM (Wimber et al., 2015). The second mechanism might be contextual binding. Previous studies found that spatial, color, or temporal context affects behavioral performance, reflecting their storage even when irrelevant (Artuso & Palladino, 2011; Cai et al., 2022; Oberauer & Vockenberg, 2009). If WM establishes item-context bindings by default, context change might trigger the reactivation of an item in WM to establish its binding with the.

### The independence of attention and storage in working memory

In our study, contralateral alpha-band suppression did not differ across repetitions or contextual changes. Typically, contralateral alpha suppression is associated with the allocation of spatial attention (Woodman et al., 2022). Given the decline in the CDA index of WM storage, one would have expected a simultaneous decline in contralateral alpha suppression, supporting the view of WM as a manifestation of endogenous attention (Gazzaley & Nobre, 2012; Kiyonaga & Egner, 2013). However, the dissociation between CDA and contralateral alpha suppression implies a differentiation between attention and WM storage (Günseli et al., 2019) and demonstrates that memory reactivation does not require spatial attention.

### Eliminating alternative explanations

In the present study, we interpreted the decline in CDA as indicative of a transition to LTM (Carlisle et al., 2011; Gunseli et al., 2014ab, Grubert et al., 2016; Reinhart et al., 2014; Reinhart & Woodman, 2014; Reinhart et al., 2016; Şentürk et al., 2024). Accordingly, we took the resurgence of the CDA as a sign of reactivating an LTM back in WM (Reinhart et al., 2014; Reinhart et al., 2016; Şentürk 2024). Below, we will discuss and eliminate five alternative explanations.

The first possibility is that the decreasing CDA via target repetition reflects a decline in attention to the repeated item rather than a transition to LTM. However, this is contradicted by the absence of a decline in contralateral alpha suppression and N2pc, indices of selective attention in perception and visual WM (Eimer, 1996; Foster & Awh, 2019; Hickey et al., 2009; Woodman et al., 2022). This dissociation is consistent with our previous work showing that CDA is sensitive to storage in WM and mostly unaffected by attention, which is tracked via contralateral alpha power suppression (Günseli et al., 2019; Hakim et al., 2019; van Driel et al., 2017). Thus, diminishing attention to the memory item likely does not account for the CDA pattern. Even if the CDA reflected decreasing attention, our interpretation of a decline in active maintenance holds, as a decrease in attention coupled with an increase in accuracy would not be possible without LTM.

The second alternative is that the increase in CDA with a new context reflects increased WM activation due to the enhanced processing of the memory display rather than retrieval from LTM. However, the lack of a corresponding increase in N2pc and contralateral alpha suppression, indices of selective attention (Eimer, 1996; Foster & Awh, 2019; Hickey et al., 2009; Woodman et al., 2022) argues against stronger encoding, as the latter requires stronger attention (Boynton, 2005; Carrasco, 2018; Mangun, 1995; Posner, 1980; Reynolds & Chelazzi, 2004; Treisman & Geffen, 1967). Importantly, whether the source is stronger encoding or retrieval from LTM, WM representations were reactivated with a context change despite the item being available in LTM. Thus, our interpretation of a context-driven activation of WM holds, regardless of whether this activation comes from reencoding or retrieving the item from LTM.

The third alternative is that the CDA increase with a context change reflects increased arousal or attention to the task. However, this is unlikely for three reasons. First, task rules are abstract representations, and arousal is a global state, neither of which would be lateralized. Therefore, they cannot explain a larger CDA, which is lateralized relative to the memory item. In line with this, previous studies found the CDA to be immune to confounds from global effects like arousal or attention (Feldmann-Wüstefeld et al., 2018; Ikkai et al., 2010). Second, previous studies found that the CDA is unaffected by the difficulty of the task the item is stored for (Gunseli et al., 2014ab; Ikkai et al, 2010; Luria et al., 2016; Şentürk et al., 2014), making it reliable for assessing representation-specific activity. Third, if the CDA increase was due to increased attention to the task, this should have been evident for context changes at item repetition 1, which was not the case (also see Şentürk et al., 2024). Therefore, a global increase in attention or arousal are unlikely candidates for explaining the CDA increase with context change.

The fourth concern regarding the CDA decline as transfer to LTM might be the lack of an effect of target repetition on reaction times. However, this was neither unexpected nor crucial. First, the decline in reaction time over target repetitions is task-dependent, being strongest in effortful search tasks and absent in recognition tasks (Carlisle et al., 2011; Gunseli et al., 2014ab). In the present study, we replicated the latter. Second, accuracy improved with repeated targets, implying learning (Adam & Vogel, 2018; Olson et al., 2005). Third, our primary outcome measure, CDA, has been consistently shown to index WM load (Roy & Faubert, 2022; Adam et al., 2018; Kang & Woodman, 2014; Feldmann-Wüstefeld et al., 2018; Hakim et al., 2019; Günseli et al., 2019, Luria et al., 2016; Vogel & Machizawa, 2005; Ikkai et al., 2010; McCollough et al., 2007), thus serving as a more direct indicator of the memory representation’s status than reaction time, and substantial evidence now correlates the decline in CDA with learning (Carlisle et al., 2011; Gunseli et al., 2014ab, Grubert et al., 2016; Reinhart et al., 2014; Reinhart & Woodman, 2014; Şentürk et al., 2024). Memory performance not worsening despite weakening neural WM indices supports long-term memory taking over given that forgetting would predict declining performance.

Besides the clear decrease in CDA with repetition, we observed a recovery in CDA at the 6th compared to the 5th repetition. This raises the question of why participants allocate more WM resources to maintaining the same information after transferring it to LTM. However, prior research has demonstrated CDA recovery at the end of the learning curve, indicating preparedness for a new item (Gunseli et al., 2014ab; Carlisle et al., 2011; Reinhart & Woodman, 2014; Şentürk et al., 2024). In our study, the fixed number of 6 repetitions provides participants with an expectation of a new item. Thus, the CDA increase on repetition 6 likely reflects preparatory WM activation. Importantly, this observation does not invalidate our core finding: the CDA on repetition 5 differs between old and new context trials, showing that individuals rely on LTM when the same context repeats and on WM when the context switches.

## Conclusion

In summary, our findings demonstrate that contextual shifts prompt memory reactivation. Items transferred to LTM through repeated storage are reactivated in WM upon encountering a task-irrelevant context change. Given the neural costs of this reactivation and the lack of observable behavioral benefits, this suggests that context-dependent reactivation is involuntary. Accordingly, we propose that memory reactivation in response to environmental changes occurs automatically, even when the context change is not directly related to the task at hand.

## Acknowledgements.

This work was funded by the Scientific and Technological Council of Türkiye (TÜBİTAK) grant (#122K700) awarded to Eren Günseli. We thank Azra Duru Erdem, Hande Altunbaş, & Öykü Özdemir for their contributions to data collection.

## Data Availability

All data and materials are shared publicly on Open Science (https://osf.io/xhrb9/?view_only=6ce0380db484406ea2070d8dbbf6ba93). Data is shared anonymously without any risk of lack of privacy.

## Code Availability

All codes of the study are shared publicly on Open Science (https://osf.io/xhrb9/?view_only=6ce0380db484406ea2070d8dbbf6ba93).

## Author Contributions

All authors contributed to the conceptualizing and the design processes of the study. Ş.Ö., Y.Ş., N.Ü. and E.G. prepared the experiment. All authors wrote the manuscript. Ş.Ö. collected the data. Ş.Ö. and E.G. conducted the data analysis. All authors discussed the interpretation of the results.

## Competing Interests

The authors declare no competing interests.

## Notes

### Competing Interest Statement

The authors have declared no competing interest.

### Summary of Updates

We shortened the introduction, added a new figure which demonstrates reaction times.

## REFERENCES

1. Adam, K. C., & Vogel, E. K. (2018). Improvements to visual working memory performance with practice and feedback. PLOS ONE, 13(8). 10.1371/journal.pone.0203279

2. Adam, K. C., Robison, M. K., & Vogel, E. K. (2018). Contralateral delay activity tracks fluctuations in working memory performance. Journal of Cognitive Neuroscience, 30(9), 1229–1240. 10.1162/jocn_a_01233

3. Alfandari, D., Belopolsky, A. V., & Olivers, C. N. (2019). Eye movements reveal learning and information-seeking in attentional template acquisition. Visual Cognition, 27(5–8), 467–486. 10.1080/13506285.2019.1636918

4. Amer, T. and Davachi, L. (2023). Extra-hippocampal contributions to pattern separation. eLife, 12: e82250. Publisher: eLife Sciences Publications, Ltd.

5. Anderson, J. R. (1983). A spreading activation theory of memory. Journal of Verbal Learning & Verbal Behavior, 22:261–295. Place: Netherlands Publisher: Elsevier Science.

6. Artuso, C. and Palladino, P. (2011). Content–context binding in verbal working memory updating: On-line and off-line effects. Acta Psychologica, 136(3):363–369.

7. Baddeley, A. (2003). Working memory: looking back and looking forward. Nature Reviews Neuroscience, 4(10):829–839. Publisher: Springer Nature.

8. Baddeley, A. (2010). Working memory. Current Biology, 20(4):R136–R140.

9. Berggren, N. and Eimer, M. (2016). Does Contralateral Delay Activity Reflect Working Memory Storage or the Current Focus of Spatial Attention within Visual Working Memory? Journal of Cognitive Neuroscience, 28(12):2003–2020.

10. Borders, A. A., Ranganath, C., and Yonelinas, A. P. (2022). The hippocampus supports high precision binding in visual working memory. Hippocampus, 32(3):217–230.eprint: https://onlinelibrary.wiley.com/doi/pdf/10.1002/hipo.23401.

11. Boynton, G. M. (2005). Attention and visual perception. Current Opinion in Neurobiology, 15(4), 465–469. 10.1016/j.conb.2005.06.009

12. Brainard, D. H. (1997). The Psychophysics Toolbox. 10:433–436.

13. Cai, Y., Fulvio, J. M., Samaha, J., and Postle, B. R. (2022). Context Binding in Visual Working Memory Is Reflected in Bilateral Event-Related Potentials, But Not in Contralateral Delay Activity. eNeuro, 9(6). Publisher: Society for Neuroscience Section: Research Article: New Research.

14. Cai, Y., Yu, Q., Sheldon, A. D., and Postle, B. R. (2018). The Role of Location-Context Binding in Nonspatial Visual Working Memory. Technical report, bioRxiv. Section: New Results Type: article.

15. Clewett, D., & Davachi, L. (2017). The ebb and flow of experience determines the temporal structure of memory. Current Opinion in Behavioral Sciences, 17, 186–193. 10.1016/j.cobeha.2017.08.013

16. Carlisle, N. B., Arita, J. T., Pardo, D., and Woodman, G. F. (2011). Attentional Templates in Visual Working Memory. Journal of Neuroscience, 31(25):9315–9322. Publisher: Society for Neuroscience Section: Articles.

17. Carrasco, M. (2018). How visual spatial attention alters perception. Cognitive Processing, 19(S1), 77–88. 10.1007/s10339-018-0883-4

18. Chota, S., Gayet, S., Kenemans, J. L., Olivers, C. N. L., and Van der Stigchel, S.(2023). A matter of availability: sharper tuning for memorized than for perceived stimulus features. Cerebral Cortex, 33(12):7608–7618.

19. Cohen, A. O., Matese, N. G., Filimontseva, A., Shen, X., Shi, T. C., Livne, E., and Hartley, C. A. (2019). Aversive learning strengthens episodic memory in both adolescents and adults. Learning & Memory, 26(7):272–279.

20. Cowan, N. (2017). The many faces of working memory and short-term storage. Psychonomic Bulletin & Review, 24(4):1158–1170.

21. Deffler, S. A., Brown, A. S., and Marsh, E. J. (2015). Judging the familiarity of strangers: does the context matter? Psychonomic Bulletin & Review, 22(4):1041–1047.

22. Draschkow, D., Kallmayer, M., & Nobre, A. C. (2021). When natural behavior engages working memory. Current Biology, 31(4). 10.1016/j.cub.2020.11.013

23. de Jong, J., Wilhelm, S., & Akyürek, E. G. (2024). Adaptive forgetting speed in working memory. Psychonomic Bulletin & Review. 10.3758/s13423-024-02507-2

24. Delorme, A. and Makeig, S. (2004). EEGLAB: an open source toolbox for analysis of single trial EEG dynamics including independent component analysis. Journal of Neuroscience Methods, 134(1):9–21.

25. Dienes, Z. (2021). How to use and report Bayesian hypothesis tests. Psychology of Consciousness: Theory, Research, and Practice, 8:9–26. Place: US Publisher: Educational Publishing Foundation.

26. Eimer, M. (1996). The N2pc component as an indicator of attentional selectivity. Electroencephalography & Clinical Neurophysiology, 99(3):225–234. Place: Netherlands Publisher: Elsevier Science.

27. Engle, R. W. (2002). Working Memory Capacity as Executive Attention. Current Directions in Psychological Science, 11(1):19–23. Publisher: SAGE Publications Inc.

28. Favila, S. E., Lee, H., and Kuhl, B. A. (2020). Transforming the Concept of Memory Reactivation. Trends in Neurosciences, 43(12):939–950.

29. Feldmann-Wüstefeld, T., Vogel, E. K., & Awh, E. (2018). Contralateral delay activity indexes working memory storage, not the current focus of spatial attention. Journal of Cognitive Neuroscience, 30(8), 1185–1196. 10.1162/jocn_a_01271

30. Foster, J. J., & Awh, E. (2019). The role of Alpha Oscillations in spatial attention: Limited evidence for a suppression account. Current Opinion in Psychology, 29, 34–40. 10.1016/j.copsyc.2018.11.001

31. Fukuda, K. and Woodman, G. F. (2017). Visual working memory buffers information retrieved from visual long-term memory. Proceedings of the National Academy of Sciences, 114(20):5306–5311. Publisher: National Academy of Sciences Section: Biological Sciences.

32. Fulvio, J. M., Yu, Q., and Postle, B. R. (2023). Strategic control of location and ordinal contextin visual working memory. Cerebral Cortex, 33(13):8821–8834.

33. Funahashi, S., Chafee, M. V., & Goldman-Rakic, P. S. (1993). Prefrontal neuronal activity in rhesus monkeys performing a delayed anti-saccade task. Nature, 365(6448), 753–756. 10.1038/365753a0

34. Fuster, J. M., & Alexander, G. E. (1971). Neuron activity related to short-term memory. Science, 173(3997), 652–654. 10.1126/science.173.3997.652

35. Gazzaley, A. and Nobre, A. C. (2012). Top-down modulation: bridging selective attention and working memory. Trends in Cognitive Sciences, 16(2):129–135.

36. Grubert, A., Carlisle, N. B., & Eimer, M. (2016). The control of single-color and multiple-color visual search by attentional templates in working memory and in long-term memory. Journal of Cognitive Neuroscience, 28(12), 1947–1963. 10.1162/jocn_a_01020

37. Godden, D. R. and Baddeley, A. D. (1975). Context-Dependent Memory in Two Natural Environments: On Land and Underwater. British Journal of Psychology, 66(3):325331. eprint:https://onlinelibrary.wiley.com/doi/pdf/10.1111/j.20448295.1975.tb01468.x.

38. Güler, B., Serin, F., & Gunseli, E. (2023). Prediction Error Is out of Context: The Dominance of Contextual Stability in Segmenting Episodic Events. 10.31234/osf.io/jgq64

39. Güler, B., Adıgüzel, Z., Uysal, B., & Günseli, E. (2024). Discrete memories of a continuous world: A working memory perspective on event segmentation. Current Research in Behavioral Sciences, 6, 100145. 10.1016/j.crbeha.2023.100145

40. Gunseli, E., Meeter, M., and Olivers, C. N. L. (2014a). Is a search template an ordinary working memory? Comparing electrophysiological markers of working memory maintenance for visual search and recognition. Neuropsychologia, 60:29–38.

41. Gunseli, E., Olivers, C. N. L., and Meeter, M. (2014b). Effects of Search Difficulty on the Selection, Maintenance, and Learning of Attentional Templates. Journal of Cognitive Neuroscience, 26(9):2042–2054.

42. Gunseli, E., Olivers, C. N., & Meeter, M. (2016). Task-irrelevant memories rapidly gain attentional control with learning. Journal of Experimental Psychology: Human Perception and Performance, 42(3), 354–362. 10.1037/xhp0000134

43. Günseli, E., Fahrenfort, J. J., van Moorselaar, D., Daoultzis, K. C., Meeter, M., and Olivers, C.N.L. (2019). EEG dynamics reveal a dissociation between storage and selective attention within working memory. Scientific Reports, 9(1):13499. Number: 1 Publisher: Nature Publishing Group.

44. Hasşerbetçi, S., & Günseli, E. (2021). EEG çalışmalarında viral Enfeksiyon Bulaşma Riskini Azaltma Yönergeleri. Türkiye Halk Sağlığı Dergisi, 19(2), 182–195. 10.20518/tjph.840203

45. Heusser, A. C., Ezzyat, Y., Shiff, I., and Davachi, L. (2018). Perceptual boundaries cause mnemonic trade-offs between local boundary processing and across-trial associative binding. Journal of Experimental Psychology: Learning, Memory, and Cognition, 44(7):1075–1090. Place: US Publisher: American Psychological Association.

46. Hickey, C., Di Lollo, V., and McDonald, J. J. (2009). Electrophysiological indices of target and distractor processing in visual search. Journal of Cognitive Neuroscience, 21(4):760–775.

47. Horner, A. J., Bisby, J. A., Wang, A., Bogus, K., and Burgess, N. (2016). The role of spatial boundaries in shaping long-term event representations. Cognition, 154:151–164.

48. Ikkai, A., Dandekar, S., and Curtis, C. E. (2016). Lateralization in Alpha-Band Oscillations Predicts the Locus and Spatial Distribution of Attention. PLoS ONE, 11(5):e0154796.

49. Ikkai, A., McCollough, A. W., and Vogel, E. K. (2010). Contralateral Delay Activity Provides a Neural Measure of the Number of Representations in Visual Working Memory. Journal of Neurophysiology, 103(4):1963–1968. Publisher: American Physiological Society.

50. Kang, M.-S., & Woodman, G. F. (2014). The neurophysiological index of visual working memory maintenance is not due to load dependent eye movements. Neuropsychologia, 56, 63–72. 10.1016/j.neuropsychologia.2013.12.028

51. Kesner, R. P. (2013). A process analysis of the CA3 subregion of the hippocampus. Frontiers in Cellular Neuroscience, 7:78.

52. Kiyonaga, A. and Egner, T. (2013). Working memory as internal attention: toward an integrative account of internal and external selection processes. Psychonomic Bulletin & Review, 20(2):228–242.

53. Konkle, T., Brady, T. F., Alvarez, G. A., and Oliva, A. (2010). Conceptual distinctiveness supports detailed visual long-term memory for real-world objects. Journal of Experimental Psychology: General, 139:558–578. Place: US Publisher: American Psychological Association.

54. Konkle, T. and Caramazza, A. (2013). Tripartite Organization of the Ventral Stream by Animacy and Object Size. Journal of Neuroscience, 33(25):10235–10242. Publisher: Society for Neuroscience Section: Articles.

55. Konkle, T. and Oliva, A. (2012). A Real-World Size Organization of Object Responses in Occipitotemporal Cortex. Neuron, 74(6):1114–1124.

56. Logan, G. D. (1988). Toward an instance theory of automatization. Psychological Review, 95:492–527. Place: US Publisher: American Psychological Association.

57. Logie, R. H., Brockmole, J. R., and Jaswal, S. (2011). Feature binding in visual short-term memory is unaffected by task-irrelevant changes of location, shape, and color. Memory & Cognition, 39(1):24–36.

58. Luria, R., Balaban, H., Awh, E., and Vogel, E. K. (2016). The contralateral delay activity as a neural measure of visual working memory. Neuroscience & Biobehavioral Reviews, 62:100–108.

59. Mangun, G. R. (1995). Neural mechanisms of visual selective attention. Psychophysiology, 32(1), 4–18. 10.1111/j.1469-8986.1995.tb03400.x

60. Marey, H. M., Semary, N. A., and Mandour, S. S. (2015). Ishihara Electronic Color Blindness Test: An Evaluation Study. Ophthalmology Research: An International Journal, pages 67–75.

61. Mızrak, E. and Oberauer, K. (2021). Working memory recruits long-term memory when it is beneficial: Evidence from the Hebb effect. Journal of Experimental Psychology: General, pages No Pagination Specified–No Pagination Specified. Place: US Publisher: American Psychological Association.

62. Nolden, S., Turan, G., Güler, B., & Günseli, E. (2024). Prediction error and event segmentation in Episodic memory. Neuroscience & Biobehavioral Reviews, 157, 105533. 10.1016/j.neubiorev.2024.105533

63. Oberauer, K. and Vockenberg, K. (2009). Updating of Working Memory: Lingering Bindings. Quarterly Journal of Experimental Psychology, 62(5):967–987. Publisher: SAGE Publications.

64. Olson, I. R., Jiang, Y., & Moore, K. S. (2005). Associative learning improves visual working memory performance. Journal of Experimental Psychology: Human Perception and Performance, 31(5), 889–900. 10.1037/0096-1523.31.5.889

65. O’Reilly, R. C. and McClelland, J. L. (1994). Hippocampal conjunctive encoding, storage, and recall: Avoiding a trade-off. Hippocampus, 4(6):661–682. _eprint: https://onlinelibrary.wiley.com/doi/pdf/10.1002/hipo.450040605.

66. Peters, B., Kaiser, J., Rahm, B., and Bledowski, C. (2021). Object-based attention prioritizes working memory contents at a theta rhythm. Journal of Experimental Psychology: General, 150:1250–1256. Place: US Publisher: American Psychological Association.

67. Posner, M. I. (1980). Orienting of attention. Quarterly Journal of Experimental Psychology, 32(1), 3–25. 10.1080/00335558008248231

68. Pu, Y., Kong, X.-Z., Ranganath, C., and Melloni, L. (2022). Event boundaries shape temporal organization of memory by resetting temporal context. Nature Communications, 13(1):622. Number: 1 Publisher: Nature Publishing Group.

69. Raccah, O., Doelling, K. B., Davachi, L., and Poeppel, D. (2022). Acoustic features drive event segmentation in speech. Journal of Experimental Psychology: Learning, Memory, and Cognition, pages No Pagination Specified–No Pagination Specified. Place: US Publisher: American Psychological Association.

70. Re, D., Inbar, M., Richter, C. G., and Landau, A. N. (2019). Feature-Based Attention Samples Stimuli Rhythmically. Current Biology, 29(4):693–699.e4.

71. Reinhart, R. M. and Woodman, G. F. (2014). High Stakes Trigger the Use of Multiple Memories to Enhance the Control of Attention. Cerebral Cortex, 24(8):2022–2035.

72. Reinhart, R. M. G., McClenahan, L. J., and Woodman, G. F. (2016). Attention’s Accelerator. Psychological Science, 27(6):790–798.

73. Reynolds, J. H., & Chelazzi, L. (2004). Attentional modulation of visual processing. Annual Review of Neuroscience, 27(1), 611–647. 10.1146/annurev.neuro.26.041002.131039

74. Robin, J., Wynn, J., and Moscovitch, M. (2016). The spatial scaffold: The effects of spatial context on memory for events. Journal of Experimental Psychology: Learning, Memory, and Cognition, 42(2):308–315. Place: US Publisher: American Psychological Association.

75. Roy, Y., & Faubert, J. (2022). Is the contralateral delay activity (CDA) a robust neural correlate for visual working memory (VWM) tasks? A reproducibility study. Psychophysiology, 60(2). 10.1111/psyp.14180

76. Sahan, M. I., Sheldon, A. D., and Postle, B. R. (2020). The Neural Consequences of Attentional Prioritization of Internal Representations in Visual Working Memory. Journal of Cognitive Neuroscience, 32(5):917–944.

77. Schönbrodt, F. D. and Wagenmakers, E.-J. (2018). Bayes factor design analysis: Planning for compelling evidence. Psychonomic Bulletin & Review, 25(1):128–142.

78. Şentürk, Y. D., Ünver, N., Demircan, C., Egner, T., & Günseli, E. (2024). The reactivation of task rules triggers the reactivation of task-relevant items. Cortex, 171, 465–480. 10.1016/j.cortex.2023.10.024

79. Sols, I., DuBrow, S., Davachi, L., & Fuentemilla, L. (2017). Event boundaries trigger rapid memory reinstatement of the prior events to promote their representation in long-term memory. Current Biology, 27(22). 10.1016/j.cub.2017.09.057

80. Stokes, M. G. (2015). ‘Activity-silent’ working memory in prefrontal cortex: a dynamic coding framework. Trends in Cognitive Sciences, 19(7):394–405.

81. Thrower, L., Dang, W., Jaffe, R. G., Sun, J. D., & Constantinidis, C. (2023). Decoding working memory information from neurons with and without persistent activity in the primate prefrontal cortex. Journal of Neurophysiology, 130(6), 1392–1402. 10.1152/jn.00290.2023

82. Treisman, A., & Geffen, G. (1967). Selective attention: Perception or response? Quarterly Journal of Experimental Psychology, 19(1), 1–17. 10.1080/14640746708400062

83. Tulving, E. and Thomson, D. M. (1973). Encoding specificity and retrieval processes in episodic memory. Psychological Review, 80(5):352–373. Place: US Publisher: American Psychological Association.

84. Vogel, E. K. and Machizawa, M. G. (2004). Neural activity predicts individual differences in visual working memory capacity. Nature, 428(6984):748–751. Number: 6984 Publisher: Nature Publishing Group.

85. Vogel, E. K., McCollough, A. W., and Machizawa, M. G. (2005). Neural measures reveal individual differences in controlling access to working memory. Nature, 438(7067):500–503. Number: 7067 Publisher: Nature Publishing Group.

86. Wang, Y. C., & Egner, T. (2022). Switching task sets creates event boundaries in memory. Cognition, 221, 104992. 10.1016/j.cognition.2021.104992

87. Wimber, M., Alink, A., Charest, I., Kriegeskorte, N., & Anderson, M. C. (2015). Retrieval induces adaptive forgetting of competing memories via cortical pattern suppression. Nature Neuroscience, 18(4), 582–589. 10.1038/nn.3973

88. Woodman, G. F., Vogel, E. K., and Luck, S. J. (2012). Flexibility in visual working memory: Accurate change detection in the face of irrelevant variations in position. Visual Cognition, 20(1):1–28. Publisher: Routledge _eprint: 10.1080/13506285.2011.630694.

89. Woodman, G. F., Wang, S., Sutterer, D. W., Reinhart, R. M. G., and Fukuda, K. (2022). Alpha suppression indexes a spotlight of visual-spatial attention that can shine on both perceptual and memory representations. Psychonomic Bulletin & Review, 29(3):681–698.

90. Xie, W. and Zhang, W. (2018). Familiarity Speeds Up Visual Short-term Memory Consolidation: Electrophysiological Evidence from Contralateral Delay Activities. Journal of Cognitive Neuroscience, 30(1):1–13.

91. Yassa, M. A. and Stark, C. E. L. (2011). Pattern separation in the hippocampus. Trends in Neurosciences, 34(10):515–525.

92. Yucel, D., Ataseven, N., Todorova, L., Güler, B., Fukuda, K., & Gunseli, E. (2023). Increased Reliance on Long-Term Memory When Anticipating Attentional Guidance. 10.31234/osf.io/cs9qa

93. Zacks, J. M. and Swallow, K. M. (2007). Event Segmentation. Current Directions in Psychological Science, 16(2):80–84. Publisher: SAGE Publications Inc.

